# Alprazolam induces anterograde amnesia for contextual fear memory and alters dorsoventral hippocampal neuronal ensembles in female mice

**DOI:** 10.1101/2025.01.08.631943

**Authors:** Kameron Kaplan, Lainey B. Toennies, Holly C. Hunsberger

**Author notes:** Corresponding Author: Holly Christian Hunsberger, Ph.D. Address: Rosalind Franklin University, 3333 Green Bay Road North Chicago, IL 60064.

## Abstract

Benzodiazepines (BZDs) are commonly prescribed anxiolytic drugs that act on GABAa receptors, and can result in anterograde amnesia, or the inability to form new memories. While BZDs have been used for decades, the brain regions and neuronal mechanisms responsible for this detrimental side effect are largely unknown at the systems neuroscience level. To analyze the effects of BZDs on long-term memory, activity-dependent ArcCreER^T2^ x eYFP mice were injected with Alprazolam 30 minutes prior to a 3-shock contextual fear conditioning (CFC) procedure and encoding ensembles were tagged with eYFP. Mice were re-exposed to the same context 5 days later, and retrieval cell activation was analyzed using the immediate early gene (IEG), c-Fos, allowing us to determine which brain regions undergo changes after alprazolam injection. Additionally, we address the question of whether alprazolam induces state-dependent memory by altering the timelines of injection. We found that 1) alprazolam treated male and female mice exhibit a decrease in memory retention and, 2) alprazolam treated female and male mice show a decrease in memory retention with saline injection prior to re-exposure, 3) alprazolam treated female mice exhibit increased EYFP+ (encoding) activation in the dCA1 and enhanced engram activation in the dCA3, and 4) alprazolam treated females showed less c-Fos+ activation in the vCA1. These results suggest that alprazolam induces sex-specific ensemble activation throughout the hippocampus and will help us understand the long-term memory deficits associated with BZD use.

## INTRODUCTION

Benzodiazepines (BZDs) are a class of anxiolytic drugs used to treat anxiety disorders such as panic disorder and post-traumatic stress disorder (PTSD)^1,2^. 30.6 million adults (12.6%) report BZD use, with women being twice as likely to use BZDs compared to men^3,4^. Although BZDs are beneficial to treat anxiety symptoms, these drugs come with the side effect of anterograde amnesia, or the inability to form new memories^5–7^. While the amnesic effect induced by BZDs is used to prevent patients from remembering stressful moments leading up to perioperative procedures, it is unknown whether chronic use of BZDs induces permanent memory impairments^8^. Clinical studies indicate that acute BZD administration induces anterograde amnesia in hippocampal dependent memory tasks such as object recognition and story recollection, suggesting a role for the hippocampus in BZD-induced amnesia^6,7^. Rodent studies have further demonstrated that BZD-induced anterograde amnesia is present in hippocampal dependent spatial and recognition memory paradigms such as the Morris Water Maze (MWM) and the Novel Object Recognition (NOR) tasks respectively^9,10^. Additionally, BZDs also induce amnesia in other hippocampal dependent associative fear learning paradigms such as the Passive Avoidance Task (PAT) and Contextual Fear Conditioning (CFC)^11,12^. While BZD-induced amnesia has been largely characterized on the behavioral level, previous studies have lacked the ability to visualize how acute administration of BZDs alter neuronal activation in the hippocampus during learning and memory tasks. Understanding these acute mechanisms is important as extensive use could result in chronic memory impairment or cognitive decline.

To visualize cell activation, activity dependent cell “tagging” transgenic mouse models, allow for the indelible labeling and visualization of neuronal ensembles or groups of neurons that show coordinated activity in response to learning^13,14^. Moreover, scientists utilize immediate early genes (IEGs), expressed shortly after neuronal activity, to visualize neuronal ensembles active during the retrieval of memory^13,15^. These models allow for the analysis of cells active during both learning and retrieval, underlying an engram or memory trace, considered the neural substrate of memory^16,17^. To understand how BZDs alter neuronal ensembles in the hippocampus during memory formation, we used the activity-dependent “tagging” mouse model, ArcCre^ERT^^2^ x EYFP^13^. Cells were labeled during a 3-shock CFC paradigm after an acute injection of the BZD, Alprazolam, in adult male and female mice. Mice were re-exposed to the fearful context to analyze fear memory expression and sacrificed in a time dependent manner to analyze neuronal ensembles and engrams within the hippocampus. We largely focused on analyzing subregions of the hippocampus as this region is essential for encoding contextual information that comprises a significant portion of contextual fear memory^18^. Here, we replicated previous studies in BZD-treated male rats and show that alprazolam treated male and female mice exhibit anterograde amnesia for long-term contextual fear memory^12^. Additionally, we found that this memory impairment is not state dependent, as mice in timelines created to circumnavigate the sedative effects of alprazolam continue to show memory impairments. Interestingly, we find that alprazolam treated female mice exhibit a significant increase in encoding cells in the dCA1 and engram activation in dCA3; however, there is a significant decrease in retrieval cell activation in the vCA1 suggesting that alprazolam alters multiple hippocampal subregions to impair memory. Surprisingly, we do not see differences in neuronal ensembles or engram activation in males suggesting that alprazolam may alter different brain regions to impair memory in a sex-specific manner.

## MATERIALS AND METHODS

### Mice

Mice were group housed (4-5/cage) in a 12-h light/dark cycle (lights on at 0600 hours) colony room maintained at 22°C. Mice had *ad libitum* access to food and water. All procedures were conducted in accordance with the National Institutes of Health regulations and by the Institutional Animal Care and Use Committee (IACUC) of Rosalind Franklin University of Medicine and Science (RFUMS).

Adult male and female (3 months old, n=5-10/group) ArcCreERT2 (+) (129S6/SvEv) bred with ROSA26-CAG-loxp-stop-loxp-EYFP-WPRE (-/-) (129S6/SvEv)^13^ mice were used in all experiments to permanently label cells active during the encoding period of a Contextual Fear Conditioning (CFC) paradigm. Genotyping was performed as previously described^13^. We did not include a naïve condition in our tagging experiments as literature has established that CFC induces significantly more cell tagging compared to context presentation alone^13^.

## Methods

### Contextual Fear Conditioning (CFC)

Adult male and female ArcCreERT2 mice were administered a 3-shock CFC paradigm. Alprazolam and saline (control) groups received an intraperitoneal (i.p.) injection of 4-hydroxytamoxifen (4-OHT, Sigma, St Louis, MO) (2mg/kg) 5 hours before the initial CFC training to allow for activity dependent neuronal tagging. Alprazolam or saline was then administered at a dose of 1 mg/kg 30 minutes before being exposed to the context chamber (Actimetrics, Wilmette, IL) equipped with a bright white light, grid floors, and lemon scent for 5 minutes. During the training period, mice received three (2 second) foot shocks (0.75 mA) every 60 seconds starting at minute 3. Mice were dark housed immediately after the training period for 3 days^13^. On day 5, mice were re-exposed to the training context containing the bright white light, grid floors, and lemon scent for 3 minutes with the absence of the shocks. Freezing data during the re-exposure period was collected (Freezeframe, Wilmette, IL) and analyzed as a proxy for memory expression.

### Shock Reactivity

Shock reactivity was tested in a separate cohort of saline and alprazolam treated male and female mice. Mice were injected with saline or alprazolam at a dose of 1 mg/kg 30 minutes before being exposed to the context chamber. Mice were allowed to habituate to the context chamber for 1 minute and then received 1 shock every 30 seconds (8 total) with increasing intensity (0.1 to 0.8 mA). Reaction to shock was recorded and the percentage of mice not reacting to the shock was plotted on a survival graph until all mice reacted to the shock.

### Dark Housing

Dark housing procedures occurred as previously described^13^. Briefly, alprazolam and saline (control) groups received an intraperitoneal (i.p.) injection of 4-OHT (2mg/kg) 5 hours before the initial CFC training and were dark housed for 3 days after to prevent any non-specific neuronal tagging. On day 4, mice were returned to their home cages to allow mice to reacclimate to a normal circadian cycle before behavioral testing on day 5. Only mice included in tagging experiments were dark housed.

### State-Dependent CFC

CFC was administered as previously described with alterations to the timeline of alprazolam and saline injections. Saline and alprazolam treated male and female mice were trained in a 3-shock CFC paradigm as described above. Mice were then randomly assigned to control or treatment groups and distributed into groups based on the re-exposure timeline. In Timeline 1, experimental groups were injected with alprazolam at a dose of 1 mg/kg 30 minutes before both CFC training and re-exposure. In Timeline 2, experimental groups were administered alprazolam at a dose of 1 mg/kg before CFC training and 0.25 mg/kg before re-exposure. In Timeline 3, experimental groups were administered alprazolam at a dose of 1 mg/kg before CFC training and received a saline injection of 1 mg/kg before re-exposure.

### Drug preparation

#### Alprazolam 1 mg/kg

A 1 mg/kg alprazolam solution (Sigma-Aldrich, A8800) was made by dissolving 1 mg of alprazolam into 0.5 ml of dimethyl sulfoxide (DMSO) and 9.5 ml of saline for a total volume of 10 ml and working concentration of 0.1 mg/ml.

#### Alprazolam 0.25 mg/kg

A 0.25 mg/kg alprazolam solution was made by dissolving 0.25 mg of alprazolam into 0.5 ml of dimethyl sulfoxide (DMSO) and 9.5 ml of saline for a total volume of 10 ml and working concentration of 0.025 mg/ml.

#### 4-hydroxytamoxifen (4-OHT)

A 10 mg/ml solution was created by dissolving 40 mg of 4-OHT (Cayman Chemical Company, 14854) into 400 ul ethyl alcohol (EtOH) and 3.6 ml of corn oil for a total volume of 4 ml. The solution was homogenized to allow for the 4-OHT to dissolve sufficiently. Mice were administered a dose of 2 mg, 5 hours before CFC training.

### Immunohistochemistry

Mice were perfused using 50ml of 1x Phosphate Buffered Saline (PBS) and 50ml of 4% paraformaldehyde (PFA) 60 minutes after CFC re-exposure in a time dependent manner to allow for immunohistochemical staining of the immediate early gene (IEG) c-Fos. Brains were left in 4% PFA overnight and then switched to a 30% sucrose solution for 2 days. Using a vibratome, brains were sliced at a thickness of 50-micron sections. To prepare for immunohistochemical staining, the tissue was first washed in 1x PBS and then incubated in a normal donkey serum (Jackson ImmunoResearch 017-000-121, 10%). We then stained for EYFP using anti-GFP Chicken (Abcam ab13970, 1:2000) and c-Fos using anti-c-Fos Rat (Synaptic Systems 226 017, 1:5000) and incubated tissue overnight at 4 degrees Celsius. On the second day, the tissue was first washed in 1x PBS and then incubated with secondary antibodies for EYFP using Cy2 conjugated Donkey anti-Chicken IgG (Jackson ImmunoResearch 703-225-155, 0.5 mg; 1:250) and c-Fos using Alexa 647 conjugated Donkey anti-Rat IgG (Thermo Fisher A78947; 1:500). Slices were then mounted onto microscope slides using antifade mountant and the slides were sealed with clear nail polish.

### Microscopy and cell quantification

Tissue was imaged using an Echo microscope at a 10x magnification. EYFP was imaged using a FITC channel, while c-Fos was imaged using a Cy5 channel. Each slice was imaged using a z-stack and a merged channel max projection image was exported for cell counting. Cell counting was performed using ImageJ. Images were split by channel to count EYFP+ and c-Fos+ cells separately. Next, the image was run through a background subtraction with a rolling ball radius of 20 pixels and a sliding paraboloid. The image threshold was then adjusted to create a binary image allowing for subtraction of background artifacts. The “despeckle” tool was then applied three times to further subtract artifacts and cell processes that were not to be counted. The “watershed” tool was then applied to allow ImageJ to discern and separate cells that were close in proximity, allowing for a more accurate count. The “remove outliers” tool was then applied to remove any last artifacts and cell processes. A region of interest (ROI) was drawn using the “polygon” tool around desired counting regions. Finally, the “analyze particles” tool retrieved the total cell count within the ROI. Particle size for each region varied based on the artifacts left in the binary image. The “overlay” tool was selected during counting for visualization of encoding (EYFP+) and retrieval cells (c-Fos+) that were included in the total cell count. Overlapping cells positive for both EYFP and c-Fos, indicative of memory traces or engrams, were counted by hand. We focused on counting regions of the hippocampus.

### Statistical analysis

All data were analyzed using JMP statistical software. CFC training data was analyzed using Repeated Measures Two-way ANOVAs (RMANOVA). CFC re-exposure data was analyzed using Two-way ANOVAs to determine the effects of sex and condition. Cell counting data was analyzed using unpaired t-tests for each hippocampal subregion. See Supplementary Table 1 for all statistics.

## RESULTS

### Alprazolam impairs contextual fear memory in male and female mice

We first sought to determine whether male and female mice responded differently to a single dose of alprazolam during fear memory as previous studies relied heavily on males. Male and female mice were injected with alprazolam or saline (1mg/kg) 30 minutes before a 3-shock CFC paradigm (**Fig. 1A**). Five days later, mice underwent context re-exposure for 3 minutes. Alprazolam treated male and female mice froze significantly more than saline control mice during the training period (**Fig. 1B**). We attribute this freezing during training to the sedative characteristics of alprazolam^19^. During re-exposure alprazolam treated mice froze significantly less, suggesting that alprazolam blunts fear memory retention and confirms previous studies (**Fig. 1C**). We have also confirmed that alprazolam-treated mice experience shock in the same way as they respond to small shock intensity (0.3mA maximum) similar to controls (**Fig. 1D**).

**Figure 1.**
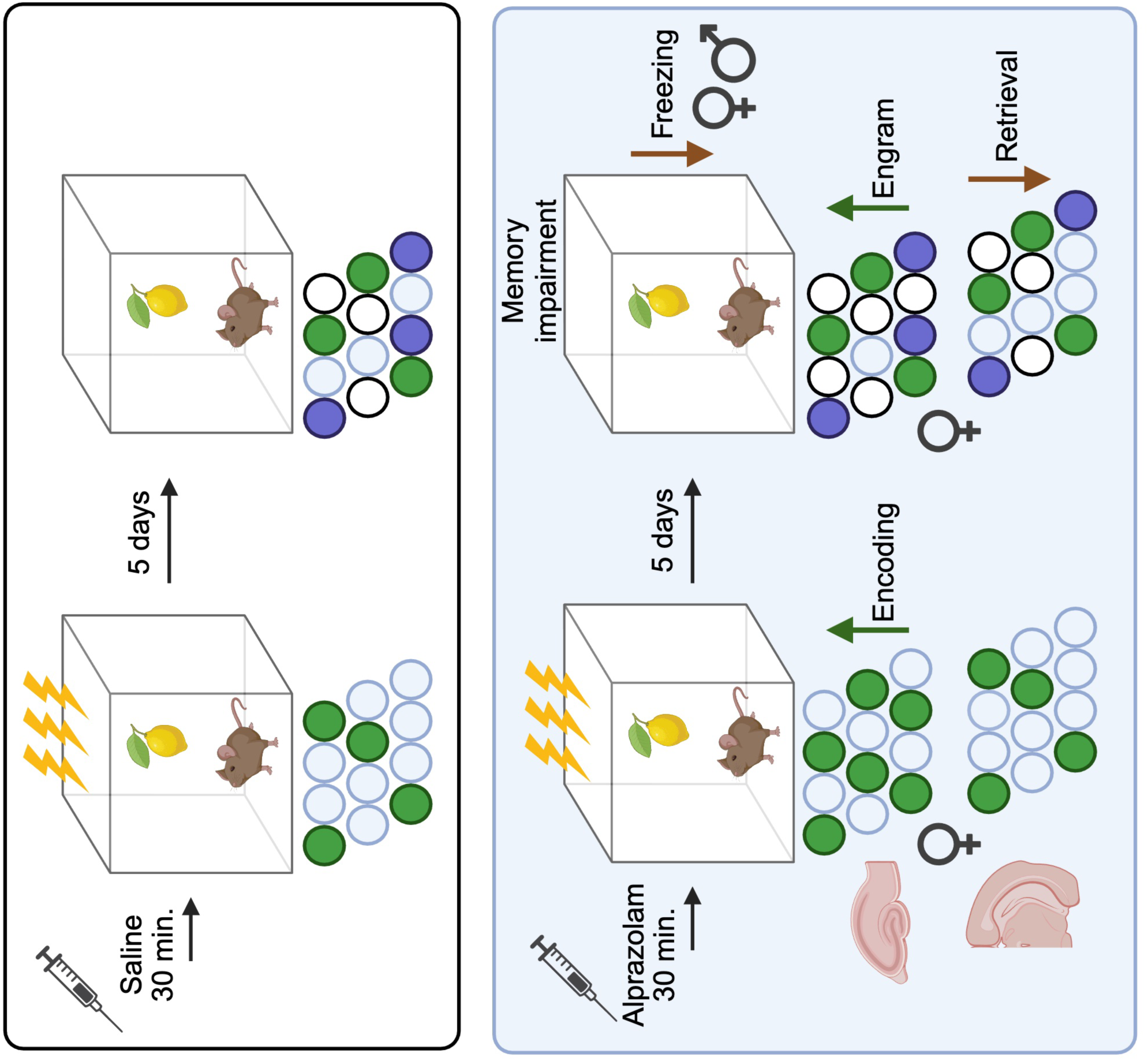
Alprazolam impairs contextual fear memory. **(A)** Contextual Fear Conditioning (CFC) experimental timeline**. (B)** Alprazolam treated male and female mice showed significantly greater freezing during the five-minute training period. (n=10-12 per group) **(C)** Alprazolam treated male and female mice showed significantly less freezing during re-exposure compared to saline treated controls. (n=10-12 per group) **(D)** Shock reactivity was comparable between male and female alprazolam and saline treated mice. (n=4-5 mice per group). Error bars represent ±SEM. *p<0.05, **p<0.01, ***p<0.001, ****p<0.0001; Alp, alprazolam; mg, milligram; kg, kilogram; sal, saline; mA, milliamp.

### Alprazolam-induced anterograde amnesia is not state dependent

To determine whether alprazolam causes state-dependent learning we created 3 alternate timelines of alprazolam injection. State-dependent learning leads to the encoding of a memory that is more easily recollected if the organism is in a similar physiological state as it was during the encoding period. Previous literature is conflicted as to whether benzodiazepines induce state dependent learning, as studies both support and contradict this hypothesis in fear memory tasks^11,20^.

Because our initial behavioral timeline did not include an injection prior to CFC re-exposure, we tested multiple timelines with injections of alprazolam or saline. Mice were assigned to saline, or alprazolam treated groups and administered a CFC training paradigm as in the previous experiment. Consistent with our previous results, alprazolam treated male and female mice froze significantly more than saline control mice during the training period over time and on average (**Fig 2A-B**). In Timeline 1, mice received either saline or alprazolam (1 mg/kg) before both CFC training and re-exposure; we did not observe any significant differences (**Fig. 2C**). However, we attribute the high freezing in alprazolam treated groups to the sedative nature of the drug. To circumnavigate the sedative properties of alprazolam during re-exposure, we developed a second timeline in which mice received saline or alprazolam (1 mg/kg) before training and 0.25 mg/kg before re-exposure. However, we did not observe any significant differences between sex or condition suggesting that even at low doses the sedative effects of alprazolam prevent movement (**Fig. 2D**). Lastly, we tested a timeline in which both alprazolam and saline treated groups were administered a saline injection before re-exposure to complete the context of receiving an injection. Timeline 3 confirmed that the alprazolam-induced memory deficit persists when groups receive a saline injection before re-exposure, suggesting that the memory deficit is not state-dependent (**Fig. 2E**).

**Figure 2.**
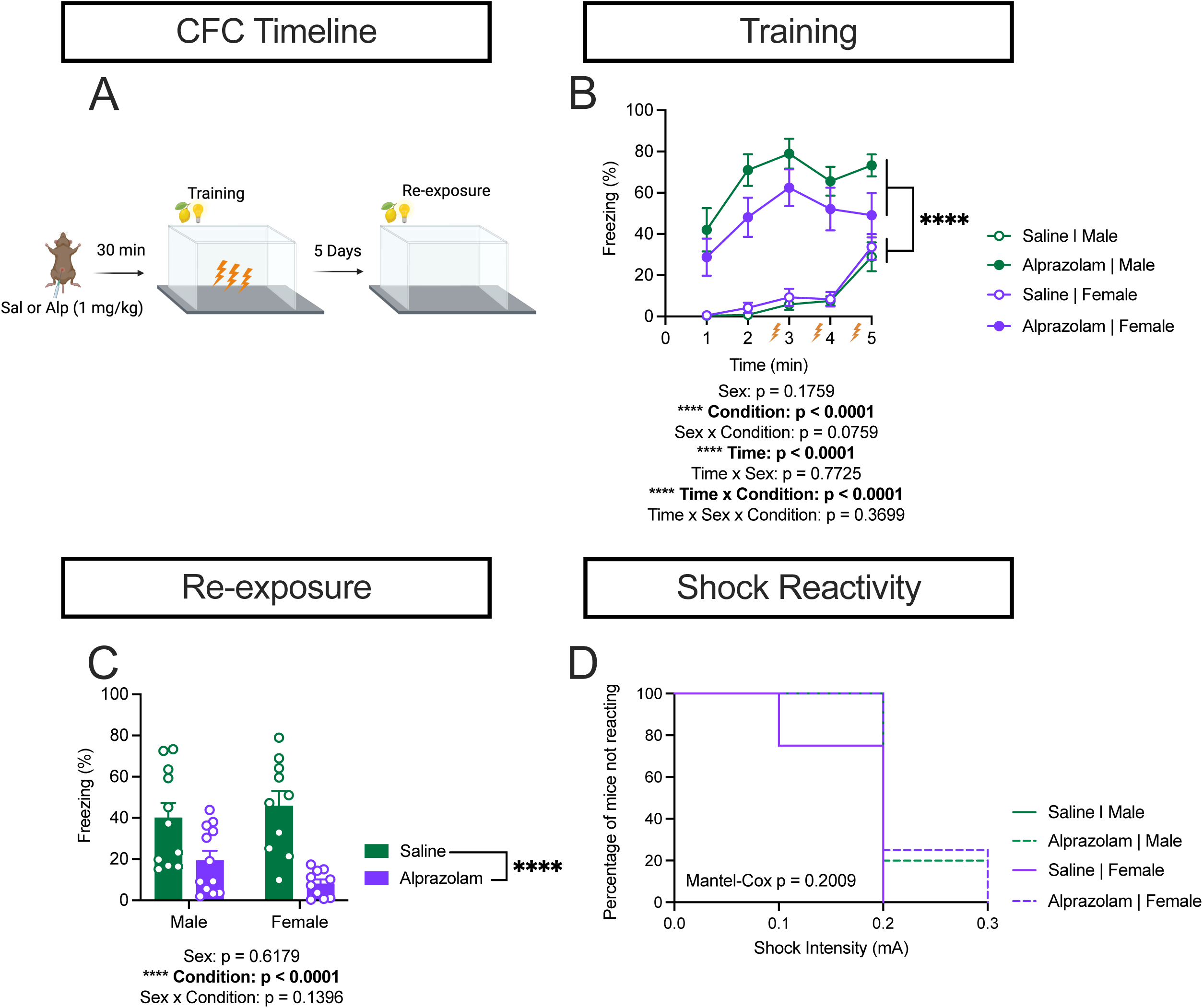
Alprazolam induced anterograde amnesia is not state dependent. **(A-B)** During CFC training, female mice showed significantly greater freezing compared to males. Alprazolam treated mice also showed significantly greater freezing compared to saline controls across time and on average. (n=15-25 per group) **(C)** Alternate timeline 1. Mice injected with saline or alprazolam (1 mg/kg) before both training and re-exposure. Freezing was comparable between alprazolam and saline treated mice during re-exposure. (n=4-9 mice per group) **(D)** Alternate timeline 2. Mice injected with saline or alprazolam (1 mg/kg) before training and saline or alprazolam (0.25 mg/kg) before re-exposure. Freezing was comparable between alprazolam and saline treated mice during re-exposure. (n=5 mice per group) **(E)** Alternate timeline 3. Mice injected with saline or alprazolam (1 mg/kg) before training and all mice received a saline injection before re-exposure. Male and female alprazolam treated mice show significantly less freezing during re-exposure. (n=5-11 mice per group). Error bars represent ±SEM. *p<0.05, **p<0.01, ***p<0.001, ****p<0.0001; min, minute; Alp, alprazolam; mg, milligram; kg, kilogram; sal, saline; CFC, contextual fear conditioning.

### Alprazolam alters neuronal ensemble and engram activation in the dorsal hippocampus of female but not male mice

To understand if alprazolam alters memory on the cellular level, we analyzed neuronal ensembles and engrams within the dorsal hippocampus in ArcCre^ERT2^ x EYFP mice after CFC. Male and female ArcCre^ERT2^ x EYFP mice were injected with 4-OHT 5 hours before CFC training to allow for Cre dependent genetic recombination leading to the indelible tagging of neurons active during the encoding period (EYFP^+^) (**Fig. 3A**). Mice were then injected with alprazolam or saline (1mg/kg) and administered a 3-shock CFC training paradigm 30 minutes later. Mice were dark housed for 3 days after CFC training to prevent any non-specific neuronal tagging. On day five, mice were re-exposed to the training context and sacrificed 60 minutes later to allow for expression of the immediate early gene (IEG), c-Fos, to analyze recent neuronal activity indicative of retrieval cells (c-Fos^+^)^13^ (**Fig. 3B**).

**Figure 3.**
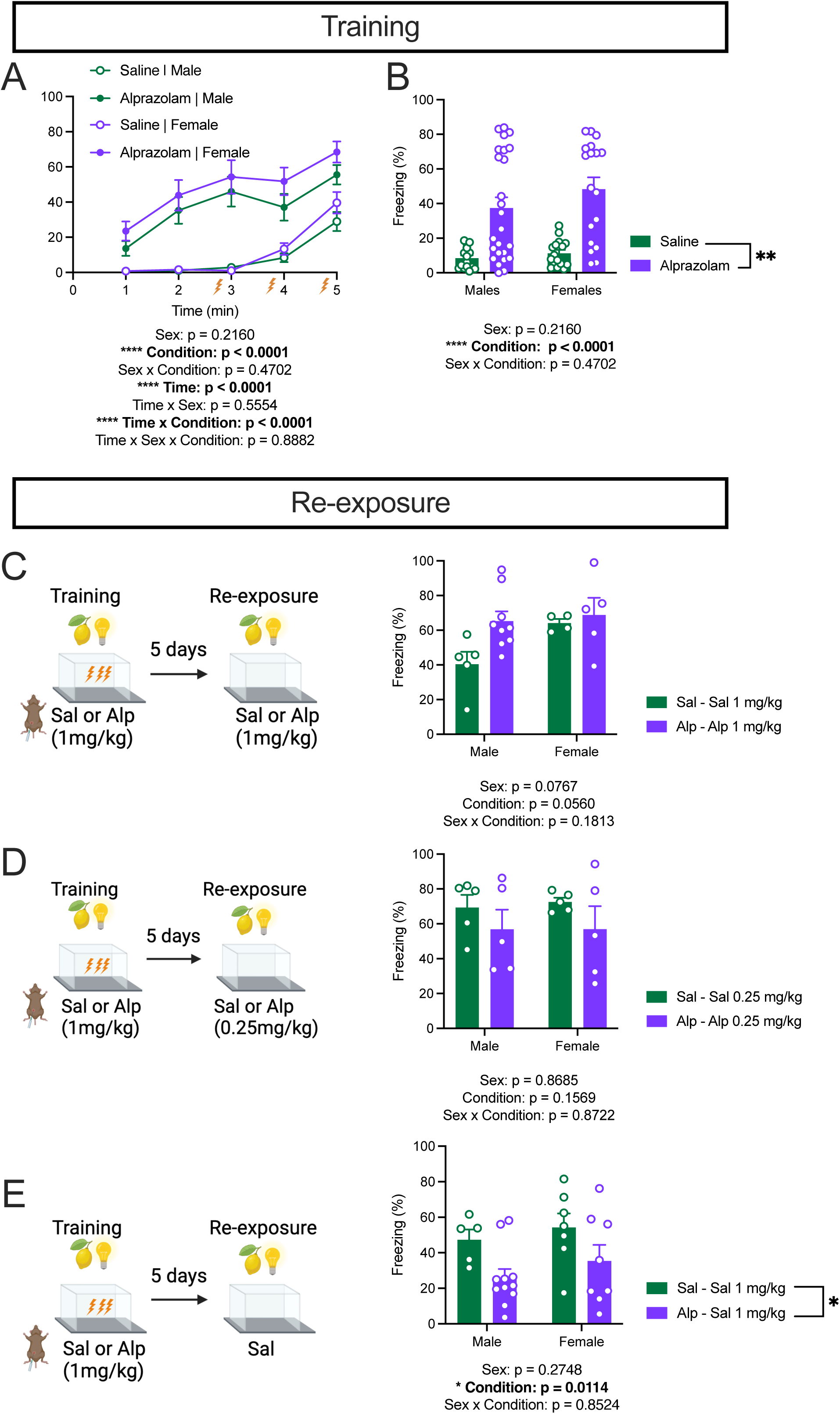
Alprazolam alters dorsal hippocampal ensembles and engrams in female but not male mice. **(A)** ArcCre^ERT2^ x EYFP genetic model and ensemble tagging. **(B)** CFC timeline for ensemble tagging. **(C)** Alprazolam treated ArcCre^ERT2^ x EYFP male and female mice show significantly less freezing during re-exposure compared to saline treated controls. **(D-E).** Alprazolam and saline treated male mice show comparable encoding and retrieval ensemble activation in the dorsal hippocampus. **(F-G)** Alprazolam and saline treated male mice show comparable engram activation when compared to encoding or retrieval ensembles. **(H-I)** Representative images of tagged ensembles in saline and alprazolam treated male mice. **(J)** Alprazolam treated female mice show significantly greater encoding ensemble activation in the dorsal CA1 of the hippocampus. **(K)** Alprazolam and saline treated female mice show comparable retrieval ensemble activation in the dorsal hippocampus. **(L)** Alprazolam and saline treated female mice show comparable engram activation in the dorsal hippocampus with overlap compared to encoding ensembles. **(M)** Alprazolam treated female mice show significantly greater engram activation in the dorsal CA3 of the hippocampus when overlap is compared to retrieval ensembles. **(N-O)** Representative images of tagged ensembles in saline and alprazolam treated female mice. White arrows indicate overlap cells. (n=5 mice per group). Error bars represent ±SEM. *p<0.05, **p<0.01, ***p<0.001, ****p<0.0001; 4-OHT, 4-hydroxytamoxifen; dDG, dorsal dentate gyrus; dCA3, dorsal cornu ammonis 3; dCA1, dorsal cornu ammonis 1; Alp, alprazolam; mg, milligram; kg, kilogram; sal, saline; CFC, contextual fear conditioning.

During re-exposure, we again found a significant condition difference, with alprazolam treated groups freezing less than saline treated controls (**Fig 3C**). While we see lower freezing levels in saline treated males, we do not attribute this to the dark housing procedure as multiple studies using this mouse model indicate normal freezing levels in control ArcCre^ERT2^ mice^13,21,22^. We did not observe any differences in encoding (EYFP^+^) or retrieval (c-Fos^+^) neuronal ensembles between alprazolam and saline treated males within the dorsal hippocampus, suggesting that this region of the hippocampus may not play a prominent role in alprazolam-induced amnesia in males (**Fig. 3D-E**). We also analyzed engram activation, or cells that had been active during the encoding and retrieval period, by calculating overlap cells as a percentage compared to both encoding (EYFP^+^) and retrieval cells (c-Fos^+^). In line with previous results, males did not exhibit differences in engram activation within the dorsal hippocampus when compared to encoding (EYFP^+^) or retrieval (c-Fos^+^) cells (**Fig. 3F-G**). Representative images of encoding and retrieval cell activation throughout the hippocampus of saline and alprazolam treated male mice are similar. (**Fig. 3H-I**).

In contrast, alprazolam treated females show enhanced encoding cell (EYFP^+^) expression in the dorsal CA1 (dCA1), suggesting alprazolam enhances encoding cell activation during the CFC training period (**Fig. 3J**). Saline and alprazolam treated female mice did not exhibit differences in retrieval cell (c-Fos^+^) expression in the dorsal hippocampus (**Fig. 3K**). When we analyzed engram activation in the dorsal hippocampus of females, we did not find differences in engram activation when % overlap was calculated and compared to encoding (EYFP^+^) cells; however, we did find a significant increase in engram activation in the dCA3 of alprazolam treated mice when % overlap was compared to retrieval (c-Fos^+^) cells, suggesting the engram is distributed into a greater number of cells in this region (**Fig. 3L-M**). Representative images of saline and Alprazolam treated female mice reveal enhanced dCA1 EYFP activation (**Fig. 3N-O**).

### Alprazolam alters neuronal ensemble activation in the ventral hippocampus of female but not male mice

We next analyzed neuronal ensemble and engram activation in the ventral hippocampus of alprazolam treated ArcCre^ERT2^ x EYFP mice after CFC. When we analyzed neuronal ensembles in male alprazolam treated mice, we did not find any differences in encoding (EYFP^+^), or retrieval (c-Fos^+^) cell activation compared to controls (**Fig. 4A-B**). We also did not find any differences in engram activation in alprazolam treated males suggesting that the ventral hippocampus does not play a role in alprazolam-induced amnesia in male mice (**Fig. 4C-D**). Representative images of tagged neuronal ensembles in the ventral hippocampus are similar in male mice (**Fig. 4E-F**).

**Figure 4.**
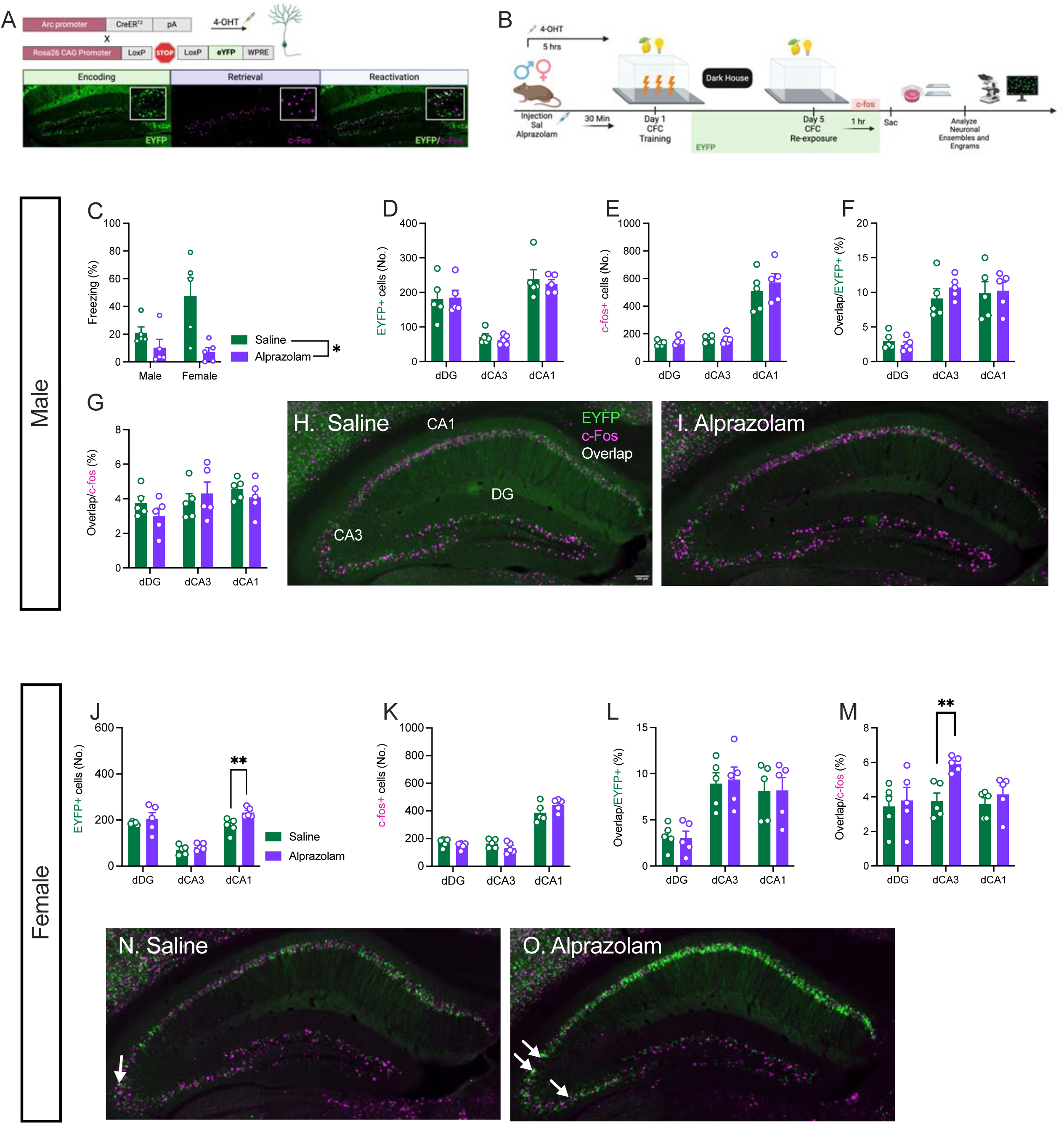
Alprazolam alters ventral hippocampal ensembles in female but not male mice. **(A-B)** Alprazolam and saline treated male mice show comparable encoding and retrieval ensemble activation in the ventral hippocampus compared to saline treated controls. **(C-D)** Alprazolam and saline treated male mice show comparable engram activation in the ventral hippocampus compared to saline controls**. (E-F)** Representative images of tagged ensembles in saline and alprazolam treated male mice**. (G)** Alprazolam and saline treated female mice show comparable encoding ensemble activation in the ventral hippocampus. **(H)** Alprazolam treated female mice show significantly less retrieval ensemble activation in the ventral CA1 compared to saline treated controls**. (I-J)** Alprazolam treated females do not show differences in engram activation in the ventral hippocampus. (n=5 mice per group). Error bars represent ±SEM. *p<0.05, **p<0.01, ***p<0.001, ****p<0.0001; vDG, ventral dentate gyrus; vCA3, ventral cornu ammonis 3; vCA1, ventral cornu ammonis 1.

When we analyzed neuronal ensembles in the ventral hippocampus of alprazolam treated female mice, we did not observe a difference in encoding (EYFP^+^) cell activation, however; we did see a significant decrease in retrieval cell (c-Fos^+^) activation in the vCA1, suggesting that the alprazolam induced memory impairment may be due to the inability to retrieve part of the memory from this brain region (**Fig. 4G-H**). Surprisingly, we did not observe any differences in engram activation in the ventral hippocampus of alprazolam treated female mice, suggesting that alprazolam did not alter the allocation of the engram in this region (**Fig. 4I-J**). Representative images reveal decreased EYFP^+^ activation in Alprazolam treated females (**Fig. 4K-L**).

## DISCUSSION

Here we have shown that alprazolam (1mg/kg) impairs contextual fear memory in adult male and female mice and that alprazolam significantly increases encoding (EYFP^+^) ensembles in the dorsal CA1 of female mice. We also found that alprazolam significantly decreases retrieval (c-Fos^+^) ensembles in the ventral CA1 of female mice. While we observe a fear memory impairment on the behavioral level, we surprisingly see a significant increase in engram activation in the dorsal CA3 of alprazolam treated female mice. However, we do not observe any significant differences in neuronal ensemble or engram activation in alprazolam treated male mice suggesting that there are sex-specific responses to alprazolam administration.

BZD users represent 12.6% of Americans, with overall use twice as prevalent in women compared to men^3,4^. Additionally, previous literature has indicated that female users may experience altered pharmacokinetic and pharmacodynamic properties of BZDs compared to men, evidenced by findings that suggest that BZD distribution in adipose tissue may be higher in females, which can alter drug serum levels and that oral contraceptives reduced drug clearance^23^. However, there is a lack of knowledge regarding sex differences and BZD-induced amnesia. Here we analyzed differences in alprazolam-induced amnesia in male and female mice at the behavioral and cellular level. Alprazolam treated male and female mice froze significantly more than controls; however, we attribute this freezing to the sedative nature of BZDs^12,19^ and confirm that these mice are experiencing the shock with our shock reactivity analysis. Others have also shown that BZD-treated rodents experience shocks during contextual fear conditioning when given moderate doses of the BZD, diazepam^12^. During re-exposure, both male and female mice experience alprazolam induced amnesia for contextual fear memory. Our data is consistent with most studies indicating the presence of BZD-induced amnesia for male rodents, and with some studies that include female rodents^9,10,12,24–27^. However, other studies indicate that gonadally intact females do not show BZD-induced amnesia for certain memory related tasks such as the PAT^28^. These inconsistencies could be due to the use of a different BZD (Lorazepam) which has different potency and kinetics compared to alprazolam and the low dose of lorazepam used in the previous studies (0.375 mg/kg)^28^. We consistently find that alprazolam (1 mg/kg) impairs contextual fear memory in male and female mice as shown in our initial behavior timeline and tagging timeline.

We next examined the state-dependency of alprazolam-induced amnesia. During training we observed the same sedative effects of alprazolam^19^. During re-exposure in Timeline 1 and 2, we do not observe alprazolam induced amnesia; however, with alprazolam in the system, we still attribute high freezing to the sedative characteristics of the drug^19^. The mice in Timeline 3, in which alprazolam-treated mice receive a saline injection to complete the context before re-exposure, continue to show alprazolam-induced memory deficits, suggesting that alprazolam-induced amnesia is not state dependent. While some literature has reported that BZD-induced amnesia is state dependent for memory paradigms such as the PAT^20^ most studies conclude that BZDs do not induce state-dependent learning for multiple memory paradigms such as PAT and Plus Maze Discriminative Avoidance Task (PM-DAT)^11,24,29^. In consensus with the latter studies, we also conclude that alprazolam is not state-dependent based on our data.

We next examined how an alprazolam injection alters memory at the cellular level in the hippocampus during CFC, as this region is essential for memory formation^30^. Contrary to our initial hypothesis, we observed enhanced encoding (EYFP^+^) ensemble activation the dorsal CA1 of alprazolam treated females. During CFC, inhibitory circuits are highly involved in fear memory encoding^31^. Specifically, the balance of excitatory and inhibitory signals coming from forebrain and brainstem regions, such as the Medial Septum (MS) and the Nucleus Incertus (NI) respectively are essential for regulating Somatostatin (SST^+^) interneurons that modulate the activity of dCA1 pyramidal neurons^31^. Therefore, we hypothesize that the enhanced activation of dCA1 pyramidal cells is a result of alprazolam-potentiated GABAergic signals coming from the (NI), leading to inhibition of SST^+^ interneurons and disinhibition of dCA1 pyramidal neurons. Previous research supports this hypothesis, as SST^+^ ensembles in the prefrontal cortex are essential for the formation and recall of auditory fear memory, and their disruption can lead to memory impairment^32^.

Additionally, the observed increase in engram activation in the dorsal CA3 of alprazolam treated females, contradicted our expected outcome of decreased engram activation after alprazolam injection. However, research indicates that the size of the engram, or number of cells within an engram, is not correlated with memory strength^16^. Therefore, while we see increased engram cell number in alprazolam treated females, this does not indicate that a functional memory has formed, evident by the memory deficit observed at the behavioral level. However, engram strength is linked to increased synaptic connections between engram cells^33^. It is possible that alprazolam potentiated inhibitory signaling prevented synaptic strengthening between engram cells; therefore, impairing memory formation, as previous literature has indicated that BZDs such as lorazepam can impair both long term potentiation (LTP) and long-term depression (LTD) in brain regions such as the perirhinal cortex^10^. Previous research has also indicated that a homeostatic excitation/inhibition (E/I) balance is essential for proper information processing^34^. Therefore, our results showing increased encoding cell activation in the dCA1, increased engram activation in the dCA3 of alprazolam treated female mice, in addition to the increased inhibitory signaling from alprazolam suggest that the proper E/I needed for the formation of a functional memory is disrupted. Subsequently, if SST^+^ neurons are silenced because of alprazolam potentiated GABAergic signaling, this would lead to increased cellular excitability which is an essential prerequisite for engram allocation^35^. This may explain why there are significantly more engram cells in the dCA3 of alprazolam treated females. The silencing of SST^+^ interneurons during encoding and enhanced cellular excitability may lead to the increase in engrams cells in addition to the memory impairment, as SST^+^ ensembles have been shown to be essential for fear memory^32^. Consistent with these hypotheses of improper memory encoding, our retrieval ensemble data indicated the inability to retrieve memory, as we see less retrieval cells in the ventral CA1, an area playing a large role in the emotional aspects of memory^30^.

Interestingly, we do not see these same results in alprazolam treated male mice. One potential explanation for these results are findings suggesting that males and females utilize brain different regions such as the hippocampal and amygdala to different degrees for fear memory retrieval, with males utilizing the dorsal hippocampus and females utilizing the basal amygdala^36^. Because males utilize the hippocampus to a larger degree, it may not be as affected by alprazolam compared to females. In the future, we plan to analyze ensemble and engram activation in different brain regions in alprazolam treated males to understand if male brains are altered by alprazolam differently. However, this was outside of the scope of this project. We will also examine SST^+^ activation and measure estrous cycle in future studies as GABAa receptor subunits responsible for tonic inhibition are known to fluctuate in response to estrous cycle phase^37^.

To conclude, while both sexes experience alprazolam induced amnesia, male and female brains are differentially altered after drug administration during contextual fear conditioning. This study is essential in understanding how BZDs such as alprazolam alter the brain at the cellular level and prevent sufficient memory formation. With this knowledge, clinicians may be inclined to find alternatives to combat disorders commonly treated with BZDs, especially in groups that are more susceptible alprazolam-induced side effects. Additionally, these early neuronal alterations could be long-lasting with repeated use and could people at risk for neurodegenerative disorders later in life.

## ACKNOWLEDGMENTS

The research reported in the article was supported by startup funds generously provided by Rosalind Franklin University of Medicine and Science. We thank the members of the Hunsberger laboratory, for their insightful comments on this project and manuscript, Dr. Joanna Dabrowska for her experimental advice and inspiration, Dr. Nicole Ferrara for her help with ImageJ analysis, and Eugene Dimitrov for his insight on shock reactivity experiments.

## AUTHOR CONTRIBUTIONS

KK was responsible for development of the idea and experimental timelines, performing experiments, data analysis, figures, and manuscript writing. LT was responsible for running behavior, drug administration, maintaining the animal colony, and tissue collection and extraction. HCH mentored, provided feedback on experiments, data, and manuscript.

---

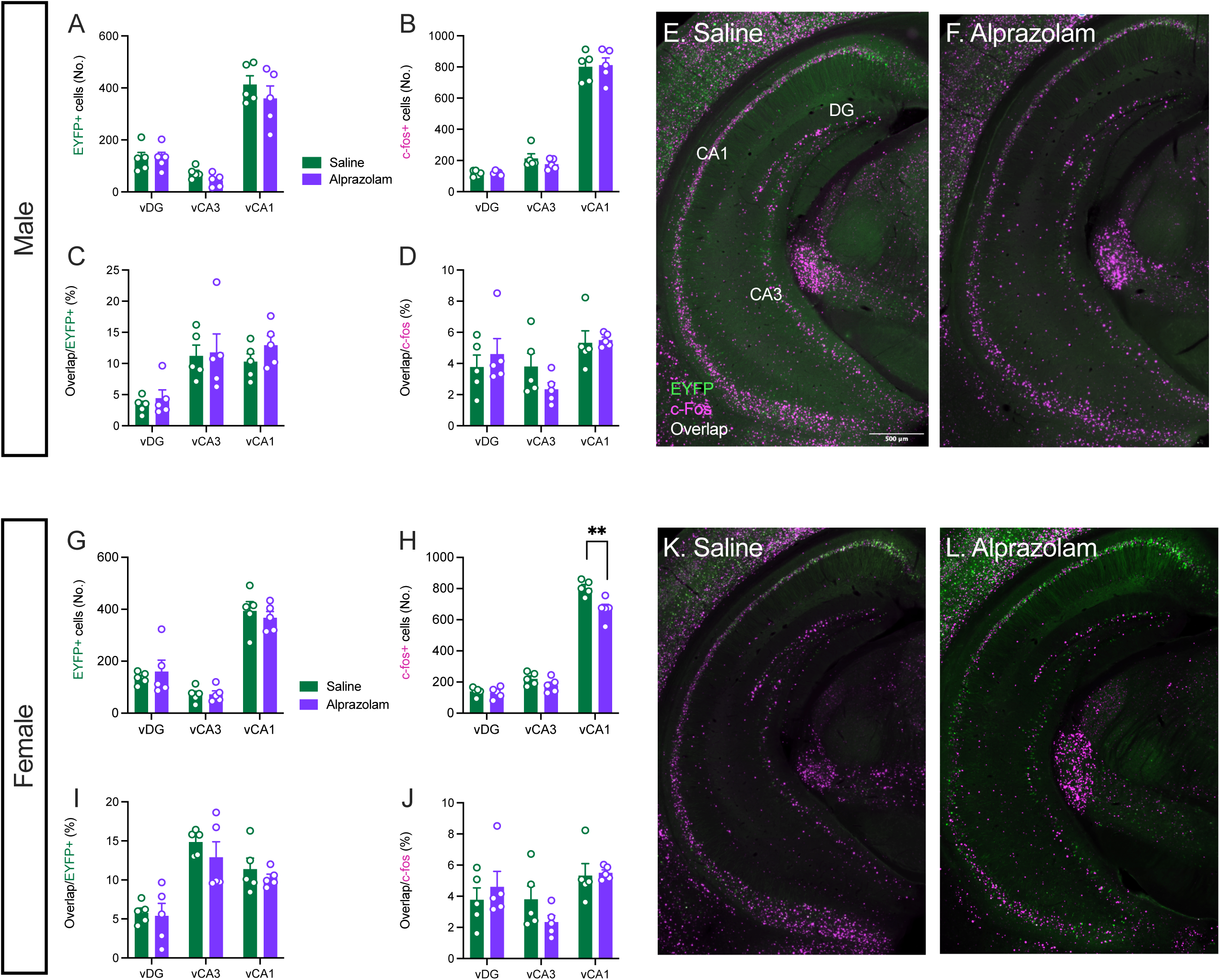

