## Supplemental for "Alprazolam induces anterograde amnesia for contextual fear memory and alters dorsoventral hippocampal neuronal ensembles in female mice"

### SUPPLEMENTARY TABLES

**Table S1.** *Statistics for mouse behavioral data.* This table lists all statistics for male and female behavioral data in **Figures 1-3**.

**Table S2.** *Statistics for cell count data.* This table lists all statistics for male and female cell count data in **Figures 3-4**.

| Table S1. Statistics for mouse behavioral data | | | | | | | | | |
|---|---|---|---|---|---|---|---|---|---|
| Cohort | Behavioral paradigm | Measurment | Statistical Test | Comparison | F | ° of freedom | p | * | Fig |
| Alprazolam before training, no alprazolam before re-exposure | CFC | Day 1 Training Freezing % | RM Two-Way ANOVA | Sex | 1.9002 | (1,39) | 0.1759 | ns | 1b |
| | | | | Condition | 69.6947 | (1,39) | <0.0001 | **** | |
| | | | | Sex x Condition | 3.3257 | (1,39) | 0.0759 | ns | |
| | | | | Time | 20.7799 | (4,36) | <0.0001 | **** | |
| | | | | Time x Sex | 0.4488 | (4,36) | 0.7725 | ns | |
| | | | | Time x Condition | 12.4491 | (4,36) | <0.0001 | **** | |
| | | | | Time x Sex x Condition | 1.1035 | (4,36) | 0.3699 | ns | |
| | | Day 2 Re-exposure Freezing % | Two-Way ANOVA | Sex | 0.2529 | (1,39) | 0.6179 | ns | 1c |
| | | | | Condition | 26.76 | (1,39) | <0.0001 | **** | |
| | | | | Sex x Condition | 2.274 | (1,39) | 0.1396 | ns | |
| Shock Reactivity | 8 shocks with increased intensity (0.1 mA - 0.8 mA) | Reaction to shock | Mantel-Cox test (Survival Curve) | - | - | 3 | 0.2009 | ns | 1d |
| State Dependent Training | CFC | Day 1 Training Freezing % | Two-Way ANOVA | Sex | 1.559 | (1,70) | 0.216 | ns | 2a |
| | | | | Condition | 35.84 | (1,70) | <0.0001 | **** | |
| | | | | Sex x Condition | 0.5272 | (1,70) | 0.4702 | ns | |
| State Dependent Training (per minute) | CFC | Day 1 Training Freezing % | RM Two-Way ANOVA | Sex | 1.559 | (1,70) | 0.216 | ns | 2b |
| | | | | Condition | 35.8443 | (1,70) | <0.0001 | **** | |
| | | | | Sex x Condition | 0.5272 | (1,70) | 0.4702 | ns | |
| | | | | Time | 59.893 | (4,67) | <0.0001 | **** | |
| | | | | Time x Sex | 0.7593 | (4,67) | 0.5554 | ns | |
| | | | | Time x Condition | 9.7011 | (4,67) | <0.0001 | **** | |
| | | | | Time x Sex x Condition | 0.2827 | (4,67) | 0.8882 | ns | |
| State Dependent (Timeline 1) re-exposure | CFC | Day 2 Freezing % | Two-Way ANOVA | Sex | 3.503 | (1,19) | 0.0767 | ns | 2c |
| | | | | Condition | 4.143 | (1,19) | 0.056 | ns | |
| | | | | Sex x Condition | 1.926 | (1,19) | 0.1813 | ns | |
| State Dependent (Timeline 2) re-exposure | CFC | Day 2 Freezing % | Two-Way ANOVA | Sex | 0.02832 | (1,16) | 0.8685 | ns | 2d |
| | | | | Condition | 2.206 | (1,16) | 0.1569 | ns | |
| | | | | Sex x Condition | 0.0267 | (1,16) | 0.8722 | ns | |
| State Dependent (Timeline 3) re-exposure | CFC | Day 2 Freezing % | Two-Way ANOVA | Sex | 1.243 | (1,27) | 0.2748 | ns | 2e |
| | | | | Condition | 7.372 | (1,27) | 0.0114 | * | |
| | | | | Sex x Condition | 0.03529 | (1,27) | 0.8524 | ns | |
| Cell Tagging cohort Re-exposure | CFC | Day 2 Freezing % | Two-Way ANOVA | Sex | 2.375 | (1,16) | 0.1428 | ns | 3c |
| | | | | Condition | 11.45 | (1,16) | 0.0038 | ** | |
| | | | | Sex x Condition | 3.851 | (1,16) | 0.0674 | ns | |

| Table S2. Statistics for cell count data | | | | | | | | |
|---|---|---|---|---|---|---|---|---|
| Statistical Test | Comparison | Sex | Measurment | Cell count | Region | p | * | Fig |
| t-test | Treatment | Males | Avg Count | EYFP | dDG | 0.9304 | - | 3d |
| | | | | | dCA3 | 0.4748 | - | |
| | | | | | dCA1 | 0.6707 | - | |
| | | | | c-Fos | dDG | 0.4526 | - | 3e |
| | | | | | dCA3 | 0.9153 | - | |
| | | | | | dCA1 | 0.4952 | - | |
| | | | % Overlap | EYFP | dDG | 0.4118 | - | 3f |
| | | | | | dCA3 | 0.3463 | - | |
| | | | | | dCA1 | 0.8576 | - | |
| | | | | c-Fos | dDG | 0.2253 | - | 3g |
| | | | | | dCA3 | 0.569 | - | |
| | | | | | dCA1 | 0.3784 | - | |
| | | Females | Avg Count | EYFP | dDG | 0.4915 | - | 3j |
| | | | | | dCA3 | 0.2662 | - | |
| | | | | | dCA1 | 0.0099 | ** | |
| | | | | c-Fos | dDG | 0.2149 | - | 3k |
| | | | | | dCA3 | 0.1911 | - | |
| | | | | | dCA1 | 0.1474 | - | |
| | | | % Overlap | EYFP | dDG | 0.8453 | - | 3l |
| | | | | | dCA3 | 0.8083 | - | |
| | | | | | dCA1 | 0.9777 | - | |
| | | | | c-Fos | dDG | 0.7337 | - | 3m |
| | | | | | dCA3 | 0.0025 | ** | |
| | | | | | dCA1 | 0.406 | - | |
| | | Males | Avg Count | EYFP | vDG | 0.9385 | - | 4a |
| | | | | | vCA3 | 0.0875 | - | |
| | | | | | vCA1 | 0.3857 | - | |
| | | | | c-Fos | vDG | 0.7438 | - | 4b |
| | | | | | vCA3 | 0.2814 | - | |
| | | | | | vCA1 | 0.8679 | - | |
| | | | % Overlap | EYFP | vDG | 0.4829 | - | 4c |
| | | | | | vCA3 | 0.8769 | - | |
| | | | | | vCA1 | 0.2168 | - | |
| | | | | c-Fos | vDG | 0.5253 | - | 4d |
| | | | | | vCA3 | 0.1634 | - | |
| | | | | | vCA1 | 0.8315 | - | |
| | | Females | Avg Count | EYFP | vDG | 0.5933 | - | 4g |
| | | | | | vCA3 | 0.9501 | - | |
| | | | | | vCA1 | 0.5572 | - | |
| | | | | c-Fos | vDG | 0.6804 | - | 4h |
| | | | | | vCA3 | 0.1723 | - | |
| | | | | | vCA1 | 0.0069 | ** | |
| | | | % Overlap | EYFP | vDG | 0.838 | - | 4i |
| | | | | | vCA3 | 0.3818 | - | |
| | | | | | vCA1 | 0.4563 | - | |
| | | | | c-Fos | vDG | 0.874 | - | 4j |
| | | | | | vCA3 | 0.8201 | - | |
| | | | | | vCA1 | 0.5934 | - | |
